# Coordinated Brain-Heart Dynamics During Human NREM Sleep

**DOI:** 10.64898/2026.09.29.755384

**Authors:** Christian A. Mikutta, Miriam Planta, Thomas Koenig, Kristoffer D. Fehér, Pauline Henckaerts, Marie Angelillo, Elisabeth Hertenstein, Carlotta L. Schneider, Marzia De Lucia, Dieter Riemann, Bernd Feige, Vladyslav V. Vyazovskiy, Christoph Nissen

## Abstract

Communication between the brain and cardiovascular system is increasingly recognized as a fundamental determinant of health, yet the mechanisms coordinating neural and cardiac rhythms remain poorly understood. Sleep provides a unique physiological state for investigating neurovisceral integration because cortical slow oscillations (SOs), thalamocortical sleep spindles, and autonomic activity undergo profound reorganization during non–rapid eye movement (NREM) sleep. We analyzed overnight high-density electroencephalography and electrocardiography recordings from 30 healthy adults and quantified interactions among SOs, sleep spindles, cardiac R-peaks, heart rate variability (HRV), spectral slope, and Lempel–Ziv complexity. To assess integrated brain–heart coordination, we developed a Joint Phase–Amplitude Coupling (JPAC) metric that quantifies the corelation between SO phase–dependent modulation of spindle amplitude and cardiac timing. SO–spindle coupling showed the expected fronto-central distribution but was not significantly related to autonomic or cortical state markers. In contrast, SO–R-peak coupling and JPAC systematically tracked physiological state. Stronger JPAC was associated with higher high-frequency HRV, steeper spectral slopes, and lower neural complexity, indicating enhanced coupling during synchronized, parasympathetically dominated NREM sleep. These findings demonstrate coordinated organization of cortical, thalamocortical, and cardiac rhythms within a common oscillatory framework and identify NREM sleep as a state and potential treatment target of large-scale neurovisceral integration.

## Introduction

Adaptive physiological regulation depends on continuous communication between the brain and cardiovascular system (1). Rather than functioning as independent organs, neural and cardiovascular systems form an integrated network that coordinates autonomic regulation, arousal, cognition, emotion, and behavior (2–7). Disruption of this coordination has been implicated in a broad range of disorders, including cardiovascular disease, depression, anxiety disorders, neurodegenerative disease, and age-related cognitive decline (1–11). Consequently, understanding how neural and cardiovascular dynamics become coordinated across physiological states represents a central challenge in systems neuroscience and medicine.

Contemporary models of neurovisceral integration propose that cardiovascular regulation emerges from distributed brain networks rather than isolated autonomic reflexes (1–7). The central autonomic network (CAN) comprises interconnected cortical, limbic, hypothalamic, and brainstem structures—including the prefrontal cortex, insula, anterior cingulate cortex, amygdala, hypothalamus, and nucleus tractus solitarius—that integrate cardiovascular, endocrine, emotional, and behavioral processes (1,6,11). Within this framework, heart rate variability (HRV), particularly high-frequency HRV (HF-HRV), is interpreted as a peripheral marker of integrated brain–body regulation and adaptive autonomic control (2–7). Together, these models suggest that cardiovascular physiology is fundamentally embedded within large-scale brain dynamics and motivate investigation of the mechanisms through which neural and cardiovascular rhythms become coordinated.

Sleep provides a unique opportunity to study neurovisceral integration under conditions of minimal behavioral and environmental interference. During non–rapid eye movement (NREM) sleep, cortical activity becomes highly synchronized, producing large-scale slow oscillations (SOs) that organize alternating periods of neuronal silence and excitation across widespread cortical networks (12–14). Concurrently, sleep spindles generated within thalamocortical circuits become temporally nested within specific phases of the SO cycle and contribute to memory consolidation and systems-level communication across the sleeping brain (15–18). At the same time, autonomic activity undergoes marked reorganization, characterized by reduced heart rate, increased parasympathetic tone, and relative cardiovascular quiescence (19–22). These coordinated neural and autonomic changes have led to the view that NREM sleep represents a privileged physiological state for investigating large-scale interactions between the central and autonomic nervous systems.

Growing evidence suggests that sleep is characterized by dynamic coupling between neural and cardiovascular activity. In a comprehensive review, de Zambotti and colleagues proposed that coordinated interactions between central and autonomic nervous systems represent a fundamental feature of sleep physiology rather than independent parallel processes (21). Consistent with this framework, heartbeat-evoked cortical responses vary systematically across vigilance states and are modulated by ongoing sleep oscillations, indicating that cardiac afferent information continues to be processed by the sleeping brain (23). Studies combining electroencephalography (EEG) and autonomic measures have further identified coordinated autonomic-central events cognitive performance (24,25). Dynamic analyses indicate that brain– heart interactions fluctuate continuously across sleep stages and cannot be explained solely by global arousal changes (26,27). More recently, coordinated oscillatory relationships between slow oscillations and autonomic dynamics have been linked to sleep-dependent restoration (24–28). Together, these findings support the emerging view that sleep constitutes a state of active neurovisceral integration.

Beyond oscillatory activity, recent evidence suggests that large-scale cortical state can be quantified using complementary electrophysiological markers. The aperiodic spectral exponent, commonly referred to as the spectral slope, reflects the decay of EEG power across frequencies and has emerged as a robust marker of cortical arousal, vigilance state, and excitation–inhibition balance (29–32). During deep NREM sleep, spectral slopes become steeper, reflecting increased neuronal synchrony and reduced cortical activation (29–31). Likewise, signal complexity measures such as Lempel–Ziv complexity (LZC) decreases during deep sleep and increase during wakefulness and rapid eye movement sleep, reflecting changes in the dynamical richness of cortical activity (33–35). Together, spectral slope and LZC provide complementary indices of global cortical organization that may help identify physiological conditions supporting large-scale neurovisceral coordination.

Despite growing interest in neurocardiac communication, previous work has largely focused on pairwise interactions. Slow oscillation–spindle coupling has been extensively characterized as a marker of thalamocortical coordination (15–18), whereas studies of brain–heart interactions have primarily examined relationships between EEG activity and HRV, heartbeat-evoked responses, or cardiac timing (21–28). Whether cortical slow oscillations, thalamocortical spindle activity, and cardiovascular rhythms become coordinated within a common physiological cycle remains unknown. Demonstrating such coordination would provide evidence for systems-level neurovisceral integration extending beyond conventional pairwise coupling models and may help explain how sleep simultaneously supports neural restoration, autonomic regulation, and cardiovascular health.

To determine whether neural and cardiovascular rhythms become coordinated within a common physiological framework during sleep, we combined overnight high-density electroencephalography and electrocardiography recordings in healthy adults and quantified interactions among slow oscillations, sleep spindles, and cardiac timing. We developed a Joint Phase–Amplitude Coupling (JPAC) metric to capture integrated brain–heart coordination across the slow oscillation cycle. We find that slow oscillations simultaneously organize spindle activity and cardiac timing, producing coordinated cortical, thalamocortical, and cardiovascular dynamics that are strongest during synchronized, vagally dominated NREM sleep, indicated by increased HF-HRV, steeper slope and lower LZC. These results identify a systems-level form of neurovisceral integration during NREM sleep and suggest that cardiovascular regulation is intrinsically embedded within large-scale brain-state dynamics.

## Results

### Analytical Overview

Figure 1 summarizes the analytical workflow. Raw EEG signals were used to identify SOs and sleep spindles (Figure 1A–B). Conventional SO–spindle coupling was quantified using phase– amplitude coupling metrics (Figure 1C), whereas SO–cardiac coupling was assessed using an event phase modulation index based on R-peak occurrence across the SO cycle (Figure 1D). We then introduced a novel Joint Phase–Amplitude Coupling (JPAC) measure quantifying the similarity between SO-phase-modulated spindle amplitude and SO-phase-modulated R-peak probability (Figure 1E). Autonomic state was characterized using heart-rate variability analyses (Figure 1F–G), whereas cortical state was quantified using spectral slope and Lempel–Ziv complexity (Figure 1H–I).

**Figure 1.**
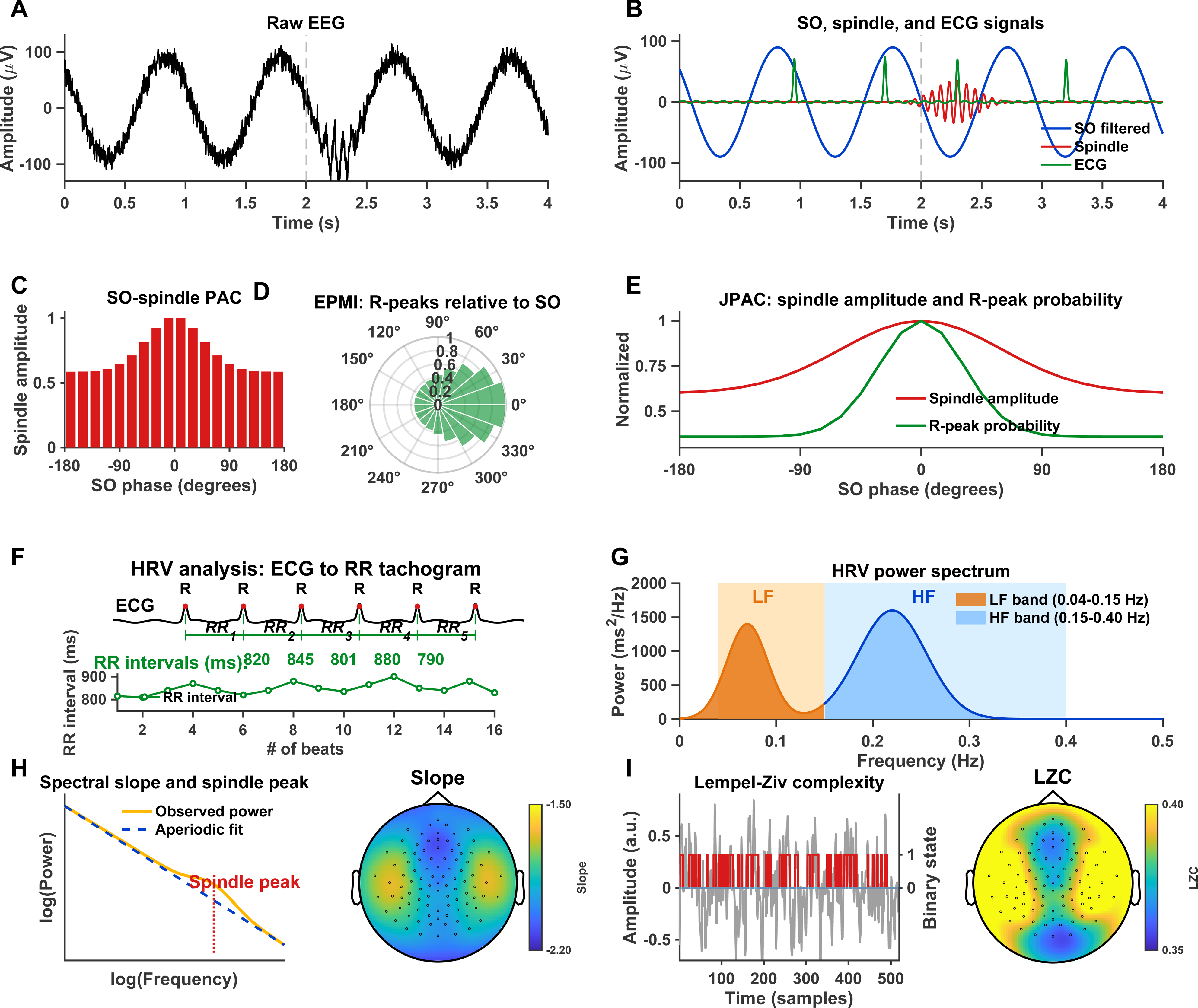
Experimental design and analysis pipeline (a) Raw EEG signal. Representative segment of the broadband EEG illustrating the presence of slow oscillatory activity with superimposed faster fluctuations. **(b) Decomposition into SO, spindle, and ECG signals.** The same segment is band-pass filtered to isolate the slow oscillation (SO; blue) and spindle-band activity (red). Spindle amplitude is shown as a continuous oscillatory envelope aligned to the SO cycle, with maximal expression during the SO upstate. Concurrently, electrocardiographic (ECG) R-peaks (green) are overlaid, revealing temporal alignment of cardiac events relative to the SO phase. **(c) SO–spindle phase–amplitude coupling (PAC).**Phase-binned spindle amplitude as a function of SO phase (−180° to 180°). Spindle amplitude is maximal around the SO upstate (0° to +90°), demonstrating canonical SO–spindle coupling. **(d) Event-phase modulation of R-peaks (EPMI).**Circular distribution of ECG R-peak occurrences relative to SO phase. R-peaks show non-uniform clustering with a preferred phase slightly preceding the SO upstate (around 0°), indicating phase-locked brain–heart interactions. **(e) Joint phase–amplitude coupling (JPAC).**Comparison of SO-phase–binned spindle amplitude (red) and SO-phase–binned R-peak probability (green), both normalized. The degree of overlap between these profiles quantifies tri-modal coordination, capturing the alignment between thalamocortical spindle dynamics and cardiac timing within the SO cycle. **(f) From ECG to HRV time series.**Electrocardiographic (ECG) recordings are used to detect R-peaks, from which successive inter-beat (RR) intervals are computed. These intervals are transformed into a continuous HRV time series, capturing beat-to-beat fluctuations in heart rate over time. **(g) HRV power spectrum (frequency domain).** Spectral decomposition of the HRV time series reveals distinct frequency components. Low-frequency (LF; 0.04–0.15 Hz) and high-frequency (HF; 0.15–0.40 Hz) bands are highlighted, representing canonical oscillatory contributions to autonomic regulation. **(H) Spectral slope analysis**. Left: Illustration of power spectral density in log–log space, showing the separation of the aperiodic (1/f-like) component from oscillatory activity. The spectral slope was estimated from the fitted aperiodic component, while spindle oscillations appear as a spectral peak above the fit. Right: Representative scalp topography of spectral slope values, reflecting the spatial distribution of the aperiodic exponent across EEG channels. **(I) Lempel–Ziv complexity (LZC).** Left: Schematic of LZC computation. The EEG signal is binarized prior to estimating signal complexity, with higher LZC indicating greater temporal irregularity and lower signal compressibility. Right: Representative scalp topography illustrating the spatial distribution of LZC across EEG channels.

### Sleep architecture and NREM spectral organization

Polysomnographic recordings confirmed physiologically normal sleep architecture in the participant group (Supplementary Table 1). Participants achieved a mean total sleep time of 452.8 ± 42.6 min with a sleep efficiency of 92.1 ± 4.8%. Sleep onset latency was 14.3 ± 10.6 min, REM sleep latency was 87.5 ± 32.4 min, and wake after sleep onset averaged 24.7 ± 18.9 min. Sleep stage distribution was within the expected range for healthy adults, comprising 7.6 ± 3.8% N1 sleep, 49.8 ± 8.2% N2 sleep, 21.4 ± 7.1% N3 sleep, and 21.2 ± 5.4% REM sleep. Arousal indices were low and consistent with consolidated sleep.

Sleep microstructure measures likewise reflected physiological NREM sleep. Participants exhibited 2956.9 ± 489.5 slow waves across the night, with a mean slow-wave amplitude of 21.5 ± 5.9 μV and a mean duration of 0.60 ± 0.01 s. The mean spectral slope was −2.5 ± 1.4 during NREM sleep and −2.5 ± 1.0 during REM sleep. Together, these findings indicate preserved sleep continuity, physiological sleep-stage distribution, and intact NREM oscillatory activity in the participant group. Detailed PSG characteristics have been reported previously by Schneider et al..The grand-average NREM power spectrum exhibited the expected 1/f-like decay across frequencies (Figure 2A). Topographical analyses revealed frequency-specific spatial patterns. Slow oscillation power was maximal over frontal regions (Figure 2B), whereas sigma power displayed a more centro-parietal distribution (Figure 2C).

**Figure 2.**
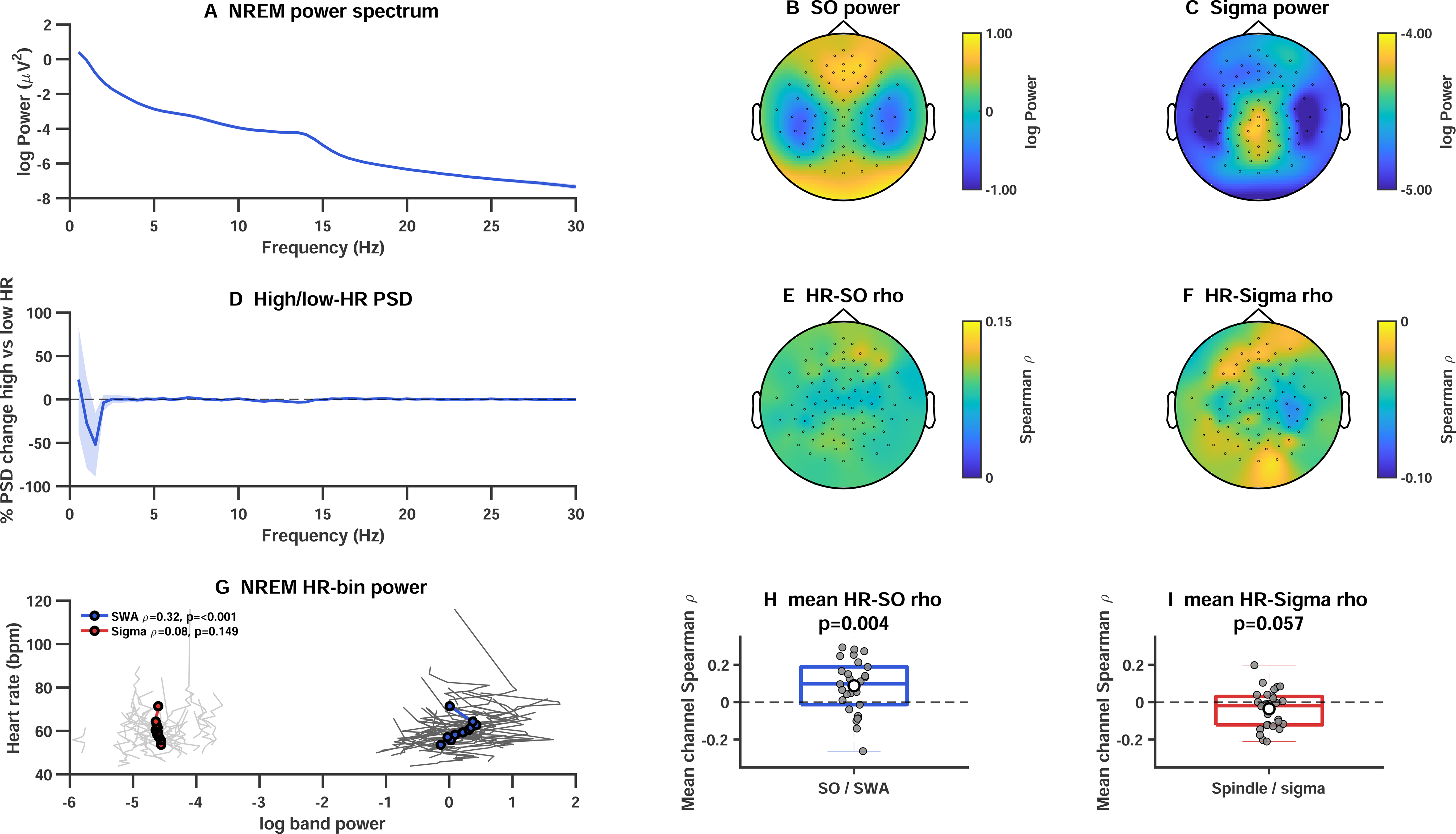
Relationship between heart rate and cortical oscillatory activity during NREM sleep. **(A)** Group-average NREM EEG power spectrum averaged across all scalp electrodes (N = 30 participants). Shaded area indicates SEM. Mean SWA (0.5–1.5 Hz) power was 0.16 ± 0.07 log(µV²), whereas mean sigma (12–16 Hz) power was −4.61 ± 0.08 log(µV²). **(B)** Topographical distribution of NREM SWA power. Slow-wave activity exhibited the expected frontal predominance. **(C)** Topographical distribution of NREM sigma power. Sigma activity showed a broadly distributed scalp pattern with relatively modest regional variation. **(D)** Mean percentage change in EEG power between the highest and lowest HR deciles. NREM epochs were divided into 10 within-participant HR quantiles and average power in the highest HR bin was compared with that in the lowest HR bin. No frequency bin survived FDR correction. Mean spectral power change across 0–30 Hz was −1.02 ± 1.06%, with the largest decrease occurring at 1.52 Hz (−52.0%) and the largest increase at 0.52 Hz (+22.8%). **(E)** Topographical distribution of the mean Spearman correlation coefficient (ρ) between HR and SWA power. Positive correlations were observed across most scalp regions, indicating that epochs with higher HR tended to exhibit greater SWA power. **(F)** Topographical distribution of the mean Spearman correlation coefficient (ρ) between HR and sigma power. Correlations were weak and predominantly negative. **(G)** Relationship between HR and oscillatory power across 10 within-participant HR quantile bins. Grey lines indicate individual participants and coloured lines indicate group means. SWA power increased significantly with HR (Spearman ρ = 0.318, p < 0.001; β = 4.10 bpm per log-power unit), whereas sigma power showed no significant association with HR (ρ = 0.083, p = 0.149). **(H)** Distribution of participant-wise mean channel HR–SWA correlations. HR–SWA correlations were significantly positive across participants (mean ρ = 0.088 ± 0.027 SEM; p = 0.0039). **(I)** Distribution of participant-wise mean channel HR–sigma correlations. HR–sigma correlations were weakly negative on average (mean ρ = −0.036 ± 0.018 SEM) and did not significantly differ from zero (p = 0.057).

### Heart rate is associated with slow-wave activity but not sigma power

To investigate relationships between cardiac dynamics and oscillatory activity, we compared spectral power between the highest and lowest HR deciles. Across the 0–30 Hz range, mean spectral power changed by −1.02 ± 1.06%, and no frequency bin survived FDR correction (Figure 2D). The largest reduction occurred at 1.52 Hz (−52.0%), whereas the largest increase occurred at 0.52 Hz (+22.8%).

Despite the absence of robust high-versus-low HR spectral differences, HR-bin analyses revealed a significant positive association between HR and SWA power. Across all HR bins, SWA increased with increasing HR (Spearman ρ = 0.318, p < 0.001; β = 4.10 bpm per log-power unit; Figure 2G). In contrast, sigma power was not significantly associated with HR (ρ = 0.083, p = 0.149; β = 2.26 bpm per log-power unit).

Consistent with these findings, channel-wise analyses demonstrated positive HR–SWA correlations across most scalp regions (Figure 2E). Participant-averaged channel correlations were significantly greater than zero (mean ρ = 0.088 ± 0.027 SEM; Wilcoxon signed-rank test, p = 0.0039; Figure 2H). The strongest association was observed over frontal electrodes, with maximal effects at Fp2 (ρ = 0.119). In contrast, HR–sigma correlations were weakly negative and did not significantly differ from zero (mean ρ = −0.036 ± 0.018 SEM; p = 0.057; Figure 2F,I).

### Robust slow oscillation–spindle coupling during NREM sleep

A non-uniform distribution of spindle events across the slow oscillation cycle was observed in 29 of 30 participants (Rayleigh test, p < 0.01). Topographical analyses of the modulation index (MI) revealed robust SO–spindle coupling with maximal effects over fronto-central scalp regions (Figure 3A). MI values showed a pronounced anterior predominance, with z-scored values exceeding 10. The topographic consistency test was significant (p = 0.001), indicating a highly reproducible spatial pattern across participants. This distribution closely matched the canonical topography of slow oscillations and sleep spindles during NREM sleep.

**Figure. 3.**
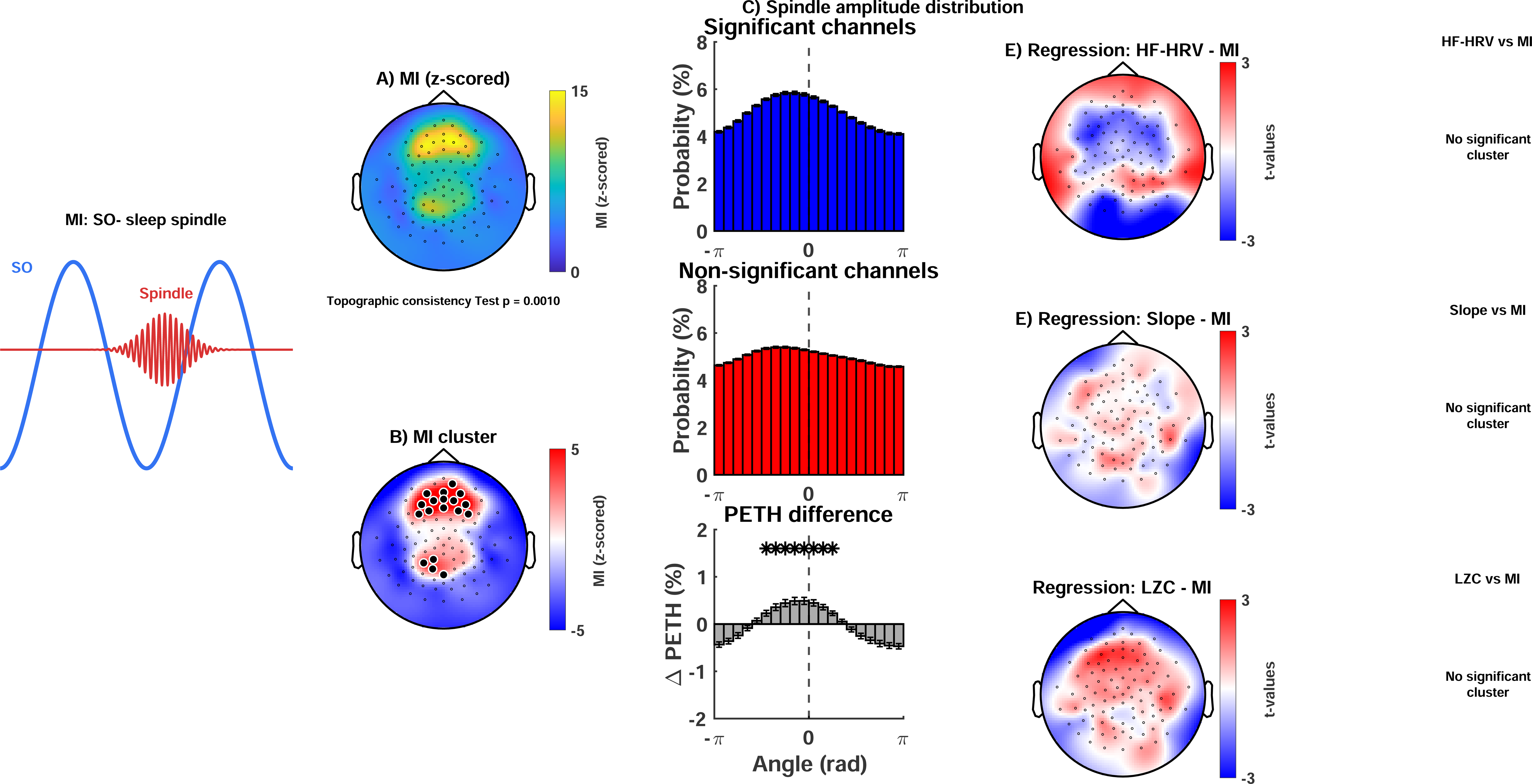
Slow oscillation–spindle coupling during NREM sleep. **(A)** Topography of SO–spindle coupling. Scalp distribution of the z-scored modulation index (MI) quantifying phase-dependent modulation of spindle amplitude by slow oscillation (SO) phase. **(B)** Group-level significance map. Topographic representation of SO–spindle coupling derived from cluster-based permutation testing. Black dots indicate electrodes belonging to significant clusters. **(C)** Phase-resolved spindle amplitude. Mean spindle-amplitude profiles across the SO cycle for electrodes within significant clusters (blue) and electrodes outside significant clusters (red). SO phase is shown from −180° to 180°. **(E)** Relationship between SO–spindle coupling and high-frequency heart rate variability (HF-HRV). Topographic distribution of regression t-values relating MI to HF-HRV across participants. **(F)** Relationship between SO–spindle coupling and spectral slope. Topographic distribution of regression t-values relating MI to the aperiodic spectral slope. **(G)** Relationship between SO–spindle coupling and Lempel–Ziv complexity (LZC). Topographic distribution of regression t-values relating MI to neural signal complexity. All regression maps depict t-values. Statistical significance was assessed using cluster-based permutation testing.

Cluster-based permutation testing confirmed that the strongest coupling effects were concentrated within a contiguous frontal cluster encompassing midline and bilateral frontal electrodes (cluster-corrected p < 0.05; Figure 3B). Phase-resolved peri-event histograms demonstrated a marked non-uniform distribution of spindle occurrence across the SO cycle (Figure 3C). Spindle probability peaked during the ascending phase and SO upstate, whereas reduced probabilities were observed around the descending phase and trough. In contrast, channels outside the significant cluster exhibited comparatively uniform phase distributions. Cluster-based analyses further identified significant phase bins surrounding the SO upstate, confirming preferential spindle occurrence during this phase of the oscillatory cycle.

We next examined whether inter-individual differences in SO–spindle coupling were related to autonomic or cortical state markers. Although exploratory regression maps suggested spatially heterogeneous associations with HF-HRV, spectral slope, and Lempel–Ziv complexity, none of these effects survived cluster-based correction for multiple comparisons (all p > 0.05; Figure 3D). Thus, unlike subsequent measures of brain–heart coupling, SO–spindle coupling did not show reliable relationships with autonomic activity or global cortical state markers. Together, these findings demonstrate robust and highly reproducible SO–spindle coupling over fronto-central regions, consistent with canonical thalamocortical coordination during NREM sleep.

### Cardiac timing is orchestarted around Slow oscillations during NREM sleep

Twenty-eight of thirty participants demonstrated significant non-uniform phase distributions of R-peaks relative to slow oscillations (Rayleigh test, p < 0.01). The z-scored event-phase modulation index (EPMI) map revealed maximal SO–R-peak coupling over fronto-central scalp regions (Figure 4A). The topographic consistency test was significant (p = 0.001), indicating a highly reproducible spatial pattern across participants.

**Figure 4.**
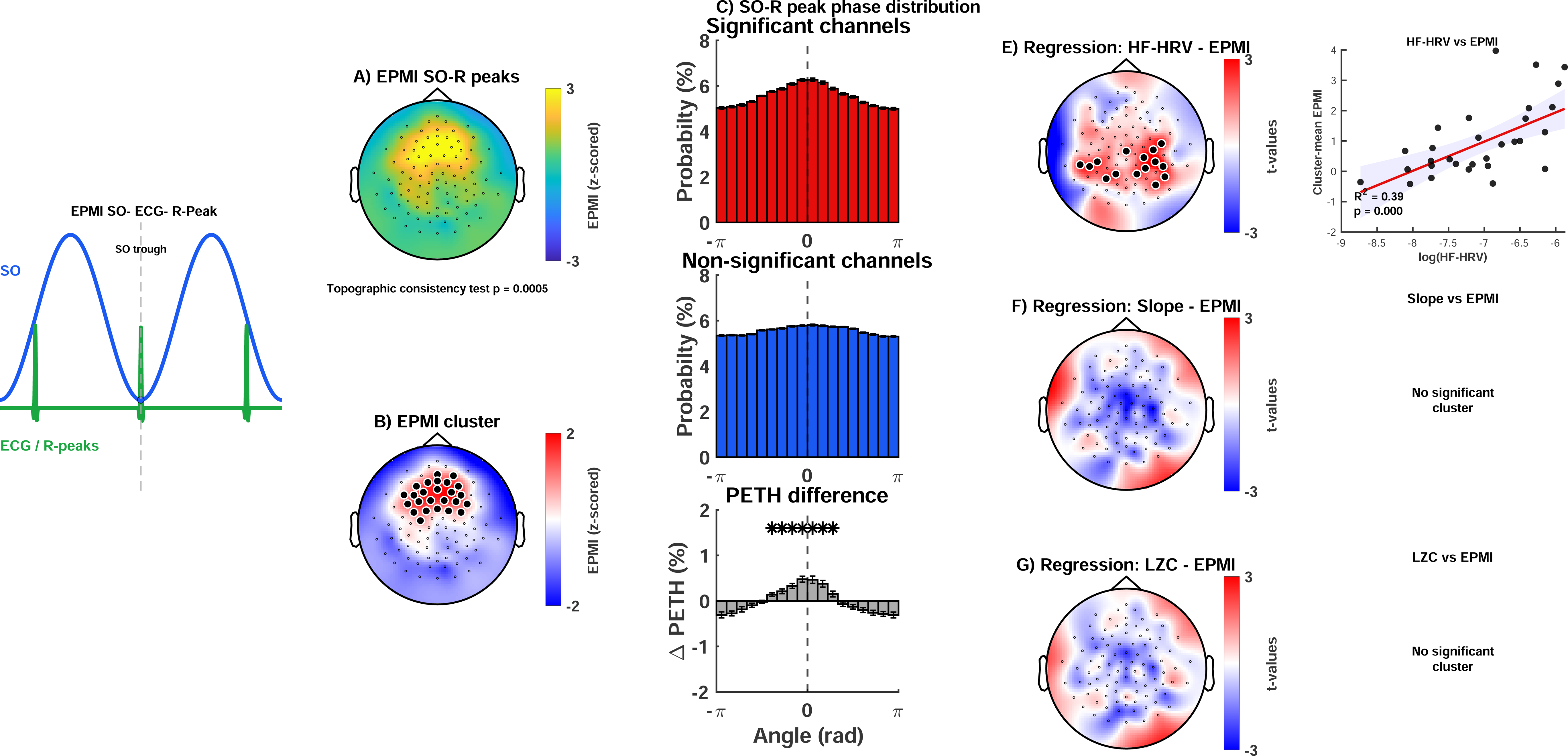
Coupling between slow oscillations and cardiac timing during NREM sleep. **(A)** Topography of SO–R-peak coupling. Scalp distribution of the z-scored Event-Phase Modulation Index (EPMI) quantifying phase-dependent modulation of ECG R-peak occurrence by slow oscillations (SOs). **(B)** Group-level significance map. Topographic representation of SO–R-peak coupling derived from cluster-based permutation testing. Black dots indicate electrodes belonging to significant clusters. **(C)** Phase-resolved R-peak probability. Mean phase-binned R-peak probability across the SO cycle for electrodes within significant clusters (blue) and electrodes outside significant clusters (red). The lower panel shows the difference between both distributions. Horizontal bars indicate phase bins reaching statistical significance following cluster-based permutation testing. **(D)** Preferred SO phase of cardiac events. Circular distributions of preferred SO phases associated with ECG R-peaks. The upper panel shows participant-wise preferred phases for electrodes within significant clusters, whereas the lower panel shows preferred phases for electrodes outside significant clusters. **(E)** Relationship between SO–R-peak coupling and high-frequency heart rate variability (HF-HRV). Topographic distribution of regression t-values relating EPMI to HF-HRV across participants. **(F)** Relationship between SO–R-peak coupling and spectral slope. Topographic distribution of regression t-values relating EPMI to the aperiodic spectral slope. **(G)** Relationship between SO–R-peak coupling and Lempel–Ziv complexity (LZC). Topographic distribution of regression t-values relating EPMI to neural signal complexity.All regression maps depict t-values. Statistical significance was assessed using cluster-based permutation testing.

Cluster-based permutation testing identified a significant fronto-central cluster in which empirical EPMI values exceeded subject-specific null distributions (cluster-corrected p < 0.05; Figure 4B). Within this cluster, peri-event phase histograms demonstrated a non-uniform distribution of R-peaks across the SO cycle, with a clear preference around the SO upstate (Figure 4C). In contrast, channels outside the significant cluster exhibited comparatively flat phase distributions. Cluster-based testing further identified significant increases in R-peak probability surrounding the SO upstate (p < 0.05).

We next examined whether SO–R-peak coupling was related to autonomic and cortical state markers. Regression analyses revealed a significant central positive cluster linking stronger SO– R-peak coupling to higher HF-HRV (cluster-corrected p < 0.05; Figure 4D), indicating enhanced coupling during states of greater parasympathetic activity. No significant associations were observed for slope and Lempel–Ziv complexity.

Together, these findings demonstrate that coupling between cortical slow oscillations and cardiac timing is spatially specific, strongest over fronto-central regions, and preferentially expressed during synchronized, parasympathetically dominated NREM sleep.

### Direct spindle–cardiac coupling is weak

In contrast to the robust coupling observed between slow oscillations and cardiac timing, direct spindle–R-peak coupling was weak. Only 3 of 30 participants demonstrated a significant non-uniform distribution of R-peaks relative to spindle events (Rayleigh test, p < 0.01). The EPMI map exhibited a heterogeneous spatial pattern with modest maxima over central regions (Supplementary Figure 1). However, cluster-based permutation testing did not identify any clusters surviving correction for multiple comparisons (all p > 0.05).

These findings indicate that cardiac timing is not systematically coupled to spindle events themselves. Rather, the stronger effects observed for SO–R-peak coupling suggest that interactions between cortical and cardiovascular rhythms are preferentially organized within the slow oscillation framework.

### Joint brain–heart coupling reveals coordinated cortical, thalamocortical, and cardiovascular dynamics

Having demonstrated robust SO–spindle coupling and SO–R-peak coupling, we next asked whether thalamocortical and cardiovascular rhythms become coordinated within the same slow oscillatory cycle. To address this question, we quantified Joint Phase–Amplitude Coupling (JPAC), a measure of the similarity between SO phase-dependent modulation of spindle amplitude and SO phase-dependent modulation of cardiac R-peak timing.

The JPAC map revealed maximal SO–spindle–R-peak coupling over fronto-central scalp regions (Figure 5A). The topographic consistency test was significant (p = 0.005), indicating a highly reproducible spatial pattern across participants. Cluster-based permutation testing identified a significant fronto-central cluster in which empirical JPAC values exceeded subject-specific null distributions (cluster-corrected p < 0.05; Figure 5B). Phase-resolved analyses demonstrated coordinated modulation of both spindle amplitude and R-peak occurrence across the SO cycle, with maxima aligned to the SO upstate (Figure 5C). Thus, cortical slow oscillations, thalamocortical spindle activity, and cardiac timing were preferentially coordinated within a common physiological cycle.

**Fig. 5.**
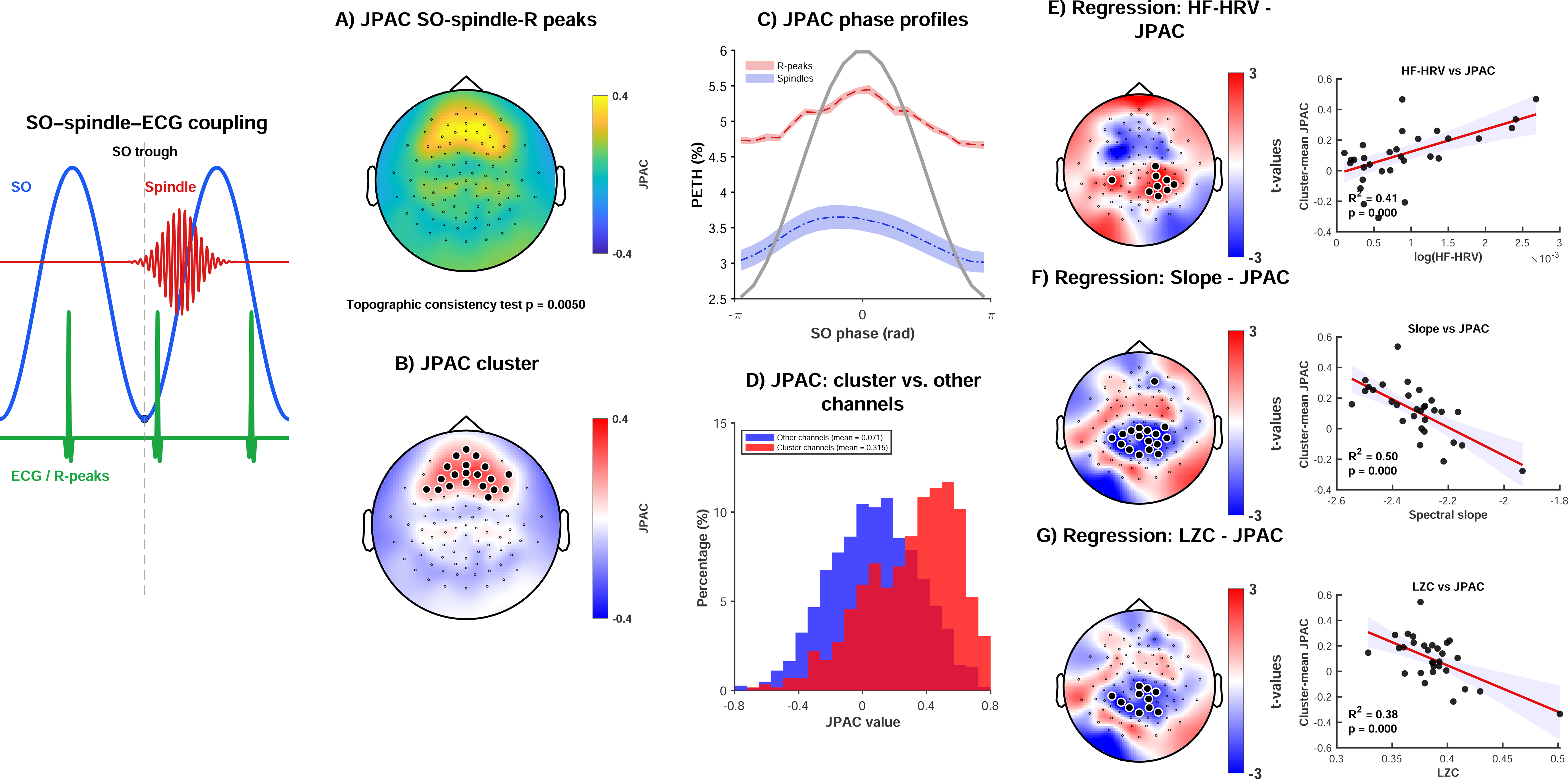
Joint phase–amplitude coupling links cortical, thalamocortical, and cardiac dynamics during NREM sleep. **(A)** Topography of Joint Phase–Amplitude Coupling (JPAC). Scalp distribution of JPAC, quantifying the correlation between the SO-phase–dependent spindle-amplitude profile and the SO-phase–dependent R-peak probability profile. The topographic consistency test was significant (p = 0.001). **(B)** Group-level significance map. Topographic representation of JPAC derived from cluster-based permutation testing. Black dots indicate electrodes belonging to significant clusters. **(C)** Phase-resolved spindle and cardiac modulation profiles. Mean spindle-amplitude profile (blue) and R-peak probability profile (red) plotted as a function of slow oscillation (SO) phase (−180° to 180°). Both measures were normalized to permit direct comparison of their phase-dependent modulation across the SO cycle. **(D)** Distribution of JPAC values. Distribution of JPAC values for electrodes within significant clusters and electrodes outside significant clusters. **(E)** Relationship between JPAC and high-frequency heart rate variability (HF-HRV). Left: topographic distribution of regression t-values relating JPAC to HF-HRV across participants. Black dots indicate electrodes belonging to significant clusters identified using cluster-based permutation testing. Right: association between cluster-averaged JPAC and HF-HRV. The red line represents the robust regression fit and the shaded region indicates the 95% confidence interval. **(F)** Relationship between JPAC and spectral slope. Left: topographic distribution of regression t-values relating JPAC to the aperiodic spectral slope. Black dots indicate electrodes belonging to significant clusters identified using cluster-based permutation testing. Right: association between cluster-averaged JPAC and spectral slope. More negative spectral slope values correspond to steeper aperiodic spectra. The red line represents the robust regression fit and the shaded region indicates the 95% confidence interval. **(G)** Relationship between JPAC and Lempel–Ziv complexity (LZC). Left: topographic distribution of regression t-values relating JPAC to neural signal complexity. Black dots indicate electrodes belonging to significant clusters identified using cluster-based permutation testing. Right: association between cluster-averaged JPAC and LZC. The red line represents the robust regression fit and the shaded region indicates the 95% confidence interval.For panels B and E–G, statistical significance was assessed using cluster-based permutation testing.

We next examined whether inter-individual differences in JPAC were related to autonomic and cortical state markers. Sensor-level regression analyses revealed a significant positive association between HF-HRV and JPAC within a right-central cluster (cluster-corrected p < 0.05; Figure 5E). Cluster-averaged JPAC increased linearly with HF-HRV (R² = 0.41, p < 0.001), indicating that stronger triple coupling was associated with enhanced parasympathetic activity.

JPAC was also significantly related to markers of global cortical state. A significant central cluster demonstrated that steeper (more negative) spectral slopes predicted stronger JPAC (R² = 0.43, p < 0.0001; Figure 5F), whereas a significant centro-parietal cluster revealed a negative relationship between Lempel–Ziv complexity and JPAC (R² = 0.41, p < 0.0001; Figure 5G). Accordingly, enhanced triple coupling emerged during cortical states characterized by increased neural synchrony and reduced signal complexity.

Together, these findings identify JPAC as the strongest and most physiologically informative coupling measure in the present study. Whereas SO–spindle coupling appeared relatively stable across physiological states and SO–R-peak coupling primarily tracked autonomic and cortical state markers, JPAC integrated both thalamocortical and cardiovascular dynamics, revealing coordinated brain–heart organization that was maximal during synchronized, parasympathetically dominated NREM sleep.

## Discussion

The present findings demonstrate that neurovisceral integration is a fundamental feature of human NREM sleep. We observed convergent evidence that cortical slow oscillations, thalamocortical spindle activity, and cardiac timing become coordinated within a common physiological cycle. While conventional SO–spindle coupling primarily reflected a stable thalamocortical timing mechanism, SO–R-peak coupling and Joint Phase–Amplitude Coupling (JPAC) systematically tracked autonomic and cortical state markers. The strongest effects emerged for JPAC, which quantified the correlation between SO phase-dependent modulation of spindle amplitude and cardiac timing. Importantly, stronger JPAC was associated with higher HF-HRV, steeper spectral slopes, and lower Lempel–Ziv complexity, indicating that multimodal brain– heart coordination is strongest during synchronized, vagally dominated NREM states. Together, these findings extend contemporary models of neurovisceral integration by demonstrating coordinated organization of cortical, thalamocortical, and cardiovascular dynamics within the same oscillatory framework (1–7, 21–28).

A central implication of these findings is that slow oscillations may serve as a temporal scaffold for large-scale physiological coordination during sleep. Slow oscillations are known to organize neuronal firing, coordinate widespread cortical communication, and temporally group spindle activity within specific phases of the oscillatory cycle (12–18). The present findings suggest that this organizational role extends beyond thalamocortical communication to include cardiovascular timing. Both spindle activity and cardiac R-peaks showed preferential alignment to the SO upstate, indicating that SOs may provide a common temporal reference through which neural and cardiovascular processes become coordinated. Rather than representing an isolated cortical rhythm, the SO may therefore act as a systems-level organizer linking multiple physiological subsystems during NREM sleep.

Consistent with previous literature, conventional SO–spindle coupling was robust, spatially organized, and highly reproducible across participants (15–18). Spindle probability peaked during the ascending phase and SO upstate, replicating established observations regarding thalamocortical synchronization and sleep-dependent information processing. Notably, however, inter-individual variability in SO–spindle coupling was not significantly associated with HF-HRV, spectral slope, or Lempel–Ziv complexity. This finding suggests that canonical SO–spindle coupling reflects a relatively stable thalamocortical timing mechanism that is largely preserved across physiological states. The absence of significant autonomic and cortical-state associations contrasts sharply with the findings for SO–R-peak coupling and JPAC and indicates that the cardiac effects observed here cannot be explained simply by stronger or weaker conventional SO–spindle coupling.

In contrast, SO–R-peak coupling demonstrated significant relationships with autonomic markers. Cardiac R-peaks occurred preferentially around the SO upstate, and coupling strength increased with higher HF-HRV and steeper spectral slopes. These observations extend previous work demonstrating heartbeat-evoked cortical responses during sleep and growing evidence for dynamic coupling between central and autonomic nervous systems during NREM sleep (21–28). Recent studies have shown that coordinated autonomic-central events contribute to cognitive performance (24,25), while dynamic analyses indicate continuous interactions between cardiac and neural rhythms across sleep stages (26,27). The present findings add a spatially specific fronto-central dimension to this literature and suggest that coupling between cortical slow oscillations and cardiac timing is strongest during highly synchronized and parasympathetically dominated sleep states.

The mechanisms underlying these effects remain speculative but are compatible with current models of the central autonomic network (CAN) and neurovisceral integration (1–7, 11, 36–40). The CAN comprises interconnected cortical, limbic, hypothalamic, and brainstem structures— including the prfrontal cortex, insula, anterior cingulate cortex, amygdala, hypothalamus, and nucleus tractus solitarius—that coordinate autonomic regulation, arousal, and interoceptive processing (1,6,11). Experimental stimulation studies demonstrate that activation of the insula and cingulate cortex can directly alter cardiac activity (36–38), whereas lesion and neuroimaging studies identify distributed autonomic control networks spanning cortical and subcortical structures (39,40). Within this framework, SO upstates may transiently engage descending autonomic pathways capable of influencing cardiac timing through sympathetic and parasympathetic projections. Conversely, cardiac afferent information transmitted through vagal and baroreceptor pathways may influence cortical excitability through brainstem–thalamocortical circuits. Although causality cannot be inferred from the present data, the observed phase relationships are consistent with bidirectional communication between cortical and cardiovascular systems.

The most important finding of the study was the identification of robust fronto-central JPAC. Unlike conventional pairwise coupling measures, JPAC quantified the coordinated modulation of spindle activity and cardiac timing by SO phase within the same physiological cycle. This observation extends previous reports of SO–spindle coordination and autonomic-central coupling (21–28) by demonstrating simultaneous organization of cortical, thalamocortical, and cardiovascular rhythms. The strong relationships between JPAC, HF-HRV, spectral slope, and Lempel–Ziv complexity suggest that multimodal brain–heart coupling is embedded within a broader physiological state characterized by enhanced parasympathetic activity, increased neural synchrony, and reduced cortical complexity. In this sense, JPAC may represent a sleep-specific systems-level indicator of neurovisceral integration, capturing coordinated regulation across multiple physiological domains rather than isolated pairwise interactions.

From a neurobiological perspective, these findings support the concept that sleep is an active state of systems-level regulation rather than a period of reduced physiological activity. Neurovisceral integration models propose that adaptive regulation emerges from coordinated interactions among distributed cortical, subcortical, autonomic, and peripheral systems (2–7). The present results extend these models by suggesting that NREM sleep provides a physiological state in which such interactions become highly organized in time. Slow oscillations may function as a common temporal framework linking cortical synchronization, thalamocortical communication, and cardiovascular regulation, thereby enabling coordinated information exchange across traditionally separate physiological domains.

The present findings also help place several emerging lines of sleep and neurocardiac research into a common framework. Previous studies have demonstrated coordinated interactions between autonomic and central nervous system activity during sleep, including heartbeat-evoked cortical responses (23), autonomic-central events linked to memory consolidation (24,25), and dynamic phase-amplitude relationships between cardiac and neural rhythms (26,27). In addition, recent work has shown that stronger SO–spindle coupling is associated with enhanced parasympathetic activity during NREM sleep (28), suggesting that thalamocortical and autonomic processes may become jointly organized. The present results extend these observations in several important ways. First, we demonstrate that slow oscillations organize both spindle activity and cardiac timing within the same physiological cycle. Second, we show that direct spindle–cardiac coupling is comparatively weak, indicating that the slow oscillation itself acts as the primary temporal scaffold for neurocardiac coordination. Third, by introducing JPAC, we provide a systems-level measure that captures simultaneous coordination among cortical, thalamocortical, and cardiovascular rhythms. Whereas previous studies primarily examined pairwise interactions between neural and autonomic signals, JPAC quantifies their integrated organization within a common oscillatory framework. These findings therefore support the view that sleep-dependent neurovisceral integration emerges from coordinated interactions among multiple physiological systems rather than isolated brain–heart relationships.

These observations may also have implications for cardiovascular health. Reduced HRV is a well-established predictor of cardiovascular morbidity and mortality across diverse populations (41–45). Contemporary concepts of the heart–brain axis increasingly emphasize that cardiovascular disease reflects not only peripheral dysfunction but also altered communication between neural and cardiovascular systems (8–11). The present findings suggest that cardiovascular recovery during sleep may depend not only on reduced heart rate or increased parasympathetic activity but also on the temporal coordination of cardiac rhythms with cortical and thalamocortical dynamics. If this interpretation is correct, disturbances of sleep-dependent neurocardiac coordination could contribute to cardiovascular vulnerability and may help explain epidemiological links between impaired sleep and cardiovascular disease.

Several limitations should be considered. First, all findings are correlational and therefore do not permit causal inference regarding the directionality of brain–heart interactions. Second, JPAC was derived from SO phase-binned profiles of spindle amplitude and R-peak probability and may be influenced by phase-estimation variability and event-selection criteria. Third, scalp EEG provides limited anatomical specificity, preventing direct identification of the neural generators underlying the observed effects. Fourth, physiological factors including respiration, baroreflex sensitivity, blood pressure dynamics, circadian influences, and sleep fragmentation were not explicitly modeled and may contribute to inter-individual variability in coupling strength (22). Finally, the study included healthy adults only, limiting conclusions regarding clinical populations. The present findings suggest that neurovisceral integration is a fundamental feature of human NREM sleep. Slow oscillations appear to provide a common temporal framework through which cortical synchronization, thalamocortical communication, and cardiovascular regulation become coordinated. By linking multimodal brain–heart coupling to parasympathetic activity, spectral slope, and neural complexity, the results identify a physiological state in which neural and cardiovascular systems operate in concert. Future studies should determine whether JPAC is altered in cardiovascular, neurological, and psychiatric disorders and whether interventions targeting sleep physiology—including closed-loop slow oscillation stimulation—can enhance neurocardiac integration and cardiovascular resilience (65).

## Materials and Methods

### Study design and participants

The present study investigated whether cortical slow oscillations (SOs), thalamocortical sleep spindles, and cardiac rhythms become coordinated within a common physiological cycle during human non–rapid eye movement (NREM) sleep. To address this question, we analysed data from 30 healthy adults, including 21 women and 9 men, with a mean age of 36±13 years, recorded during a baseline night as part of a previously published study on sleep–wake perception and sleep physiology (46). Participants were free of current psychiatric, neurological, cardiovascular, and sleep disorders and had not taken psychoactive medication within two weeks before participation. Additional inclusion criteria required stable sleep–wake schedules, consumption of less than 500 mg caffeine per day, and smoking of fewer than five cigarettes per day. Participants were excluded if screening polysomnography revealed clinically relevant sleep pathology, including obstructive sleep apnea (apnea–hypopnea index >15 events h⁻¹) or periodic limb movements associated with arousals (>10 events h⁻¹).

All participants provided written informed consent prior to participation. The study was conducted in accordance with the Declaration of Helsinki and approved by the Ethics Committee of the Canton of Bern (Project ID: 2019-02426).

### Sleep recordings

Participants underwent one adaptation night followed by an experimental baseline night in the sleep laboratory. Recordings were obtained using a 128-channel high-density EEG system (MicroCel Geodesic Sensor Net, Electrical Geodesics Inc., Eugene, OR, USA) together with ECG and standard polysomnographic channels. Signals were amplified using a Net Amps 400 amplifier system and digitized at 500 Hz.

The adaptation night included electrooculography, submental electromyography, respiratory effort belts, nasal airflow monitoring, pulse oximetry, and bilateral tibialis anterior electromyography to screen for sleep disorders. Sleep stages were visually scored according to the criteria of the American Academy of Sleep Medicine (47). Participants were allotted 8 h time in bed aligned to their habitual sleep schedule.

### EEG preprocessing

EEG preprocessing was performed in MATLAB R2023a (MathWorks, Natick, MA, USA) using EEGLAB (47) and FieldTrip (48). Signals were downsampled from 500 to 200 Hz, notch-filtered at 50 Hz (47–53 Hz), and band-pass filtered between 0.1 and 95 Hz. Data cleaning followed principles of the PREP pipeline (49), including robust average rereferencing, automated bad-channel detection, and spherical interpolation of rejected electrodes.

Residual artifacts were identified using a semi-automated procedure combining spectral screening and visual inspection (50). Channels exhibiting persistent noise or poor signal quality were excluded and interpolated using neighboring electrodes. Data were subsequently rereferenced to the average reference.

### Regression-based removal of electrocardiographic contamination from electroencephalography

Because ECG-related electrical activity can contaminate scalp EEG recordings (51–53), ECG artifacts were removed using channel-wise linear regression. For each EEG channel, variance explained by the simultaneously recorded ECG signal was estimated and subtracted, yielding ECG-corrected EEG signals used in all subsequent analyses.

### ECG preprocessing

ECG recordings were preprocessed using Kubios HRV Scientific version 4.1.1 (54). Signals were visually inspected and filtered according to standard procedures. R-peaks were detected using adaptive threshold algorithms followed by manual verification. Ectopic beats, missed detections, and noise-related artifacts were corrected using Kubios artifact-correction routines.

Artifact-corrected R-peak timings and RR intervals were exported for further analyses. Heart rate (HR) was calculated from successive RR intervals, and high-frequency heart rate variability (HF-HRV; 0.15–0.40 Hz) was derived according to established guidelines (55–57). These measures were subsequently related to indices of SO–spindle coupling, SO–R-peak coupling, and Joint Phase–Amplitude Coupling (JPAC).

### Detection of slow oscillations and sleep spindles

All event detection analyses were restricted to NREM sleep (N2 and N3). Slow oscillations (SOs) were detected from EEG signals band-pass filtered between 0.16 and 1.25 Hz. Candidate events were identified from successive zero crossings and retained if their duration was between 0.8 and 2.0 s and their trough-to-peak amplitude exceeded the channel-specific 75th percentile. These criteria follow established approaches for detecting physiologically relevant SOs in human sleep EEG and for quantifying SO–spindle interactions (15–17). To reduce variability related to absolute signal magnitude across participants and electrodes, individual SO waveforms were z-scored in the time domain before further analyses. For each event, the SO trough served as the temporal reference point and the instantaneous SO phase was obtained using the Hilbert transform.

Sleep spindles were detected from EEG signals filtered between 12 and 16 Hz. The analytic amplitude envelope was computed using the Hilbert transform, and spindle candidates were identified when the envelope exceeded two standard deviations above the channel-specific mean. Events with durations between 0.5 and 3.0 s were retained, consistent with standard spindle definitions and previous SO–spindle coupling studies (14–17). For each spindle, the time point of maximal spindle amplitude was defined as the spindle peak and used in subsequent phase-based analyses.

### SO–spindle coupling

To quantify the temporal coordination between slow oscillations and sleep spindles, the instantaneous phase of the SO signal was extracted from the band-pass filtered SO time series (0.16–1.25 Hz) using the Hilbert transform. SO phase was represented on a circular scale spanning −π to π radians.

For each detected spindle, spindle-band amplitude was assigned to the corresponding SO phase at the time of spindle occurrence. SO phases were subsequently grouped into 18 equally spaced bins (20° per bin), and mean spindle amplitude was calculated for each phase bin. This procedure yielded a phase-resolved spindle-amplitude distribution for each participant and electrode.

The strength of SO–spindle coupling was quantified using a modulation index (MI) adapted from established phase–amplitude coupling approaches (58). MI values were calculated by comparing the observed phase-dependent spindle-amplitude distribution with a uniform distribution, such that larger values indicated stronger preferential occurrence of spindle activity at specific phases of the SO cycle.

Preferred spindle phase and phase concentration were assessed using circular statistics implemented in CircStat (59). Statistical significance of phase clustering was evaluated using the Rayleigh test.

### SO–ECG R-peak coupling

To quantify coupling between cortical slow oscillations and cardiac timing, ECG R-peaks were assigned to the instantaneous phase of the nearest SO cycle. R-peak occurrences were binned into the same 18 SO-phase bins used for SO–spindle analyses, yielding phase-resolved probability distributions of cardiac events.

The strength of SO–R-peak coupling was quantified using an Event-Based Modulation Index (EPMI), analogous to the modulation index used for spindle analyses. EPMI measured the deviation of the observed R-peak phase distribution from a uniform distribution, with larger values indicating stronger phase-specific modulation of cardiac timing by the SO cycle.

To assess statistical significance, surrogate distributions were generated by randomly shifting R-peak timings relative to SO events while preserving the temporal structure of the cardiac signal. Empirical EPMI values were converted into z scores relative to the surrogate distribution. Preferred cardiac phases were assessed using circular statistics and Rayleigh tests (59).

### Joint Phase–Amplitude Coupling (JPAC)

To quantify integrated coordination among cortical slow oscillations, thalamocortical spindle activity, and cardiac timing, we developed a Joint Phase–Amplitude Coupling (JPAC) metric. JPAC was designed to assess whether spindle amplitude and cardiac R-peak probability exhibit similar modulation across the slow oscillation (SO) cycle.

For each participant and electrode, spindle amplitudes and R-peak occurrences were assigned to 18 equally spaced SO-phase bins. This yielded two phase-resolved profiles: (i) mean spindle amplitude as a function of SO phase and (ii) R-peak probability as a function of SO phase.

JPAC was defined as the Pearson correlation coefficient between these two phase-binned distributions. Positive JPAC values indicate that phases associated with increased spindle amplitude also exhibit increased R-peak probability, whereas values near zero indicate relative independence of spindle and cardiac modulation across the SO cycle. Thus, JPAC quantifies the extent to which thalamocortical spindle activity and cardiac timing are jointly organized by the underlying SO rhythm.

To assess statistical significance, surrogate distributions were generated by randomly circularly shifting the phase-binned spindle-amplitude profile relative to the phase-binned R-peak probability profile while preserving the shape of both distributions. This procedure was repeated 1,000 times for each participant and electrode. Empirical JPAC values were subsequently converted to z scores relative to the surrogate distribution and used for all group-level analyses. Positive z-scored JPAC values therefore indicate coupling strengths exceeding those expected by chance.

### Spectral slope and Lempel–Ziv complexity

To characterize global cortical state during NREM sleep, we quantified the aperiodic spectral slope and Lempel–Ziv complexity (LZC) of the EEG signal.

Spectral slope was estimated from the aperiodic component of the EEG power spectrum using FieldTrip’s FOOOF-based spectral parameterization (cfg.output = ‘fooof_aperiodic’) (30, 48). For each participant and electrode, EEG data were submitted to multitaper frequency analysis using a Hanning taper, and the aperiodic component was extracted across the predefined frequency range. The spectral slope was defined as the exponent parameter of the aperiodic fit. More negative values indicate steeper spectral decay and have been associated with reduced arousal, increased cortical synchrony, and deeper NREM sleep states (29–32).

Signal complexity was quantified using Lempel–Ziv complexity (LZC), a measure of the diversity and compressibility of neural activity patterns (33–35, 61). EEG signals were binarized relative to the median amplitude, and LZC was calculated according to the original Lempel–Ziv algorithm (61). Higher LZC values indicate greater temporal complexity and dynamical richness, whereas lower values reflect more stereotyped and synchronized neural activity. Participant-level LZC values were averaged across NREM sleep epochs and electrodes before subsequent analyses. Spectral slope and LZC were subsequently related to SO–spindle coupling, SO–R-peak coupling, and JPAC using sensor-level regression analyses.

### Heart-rate binning and spectral analyses

To investigate the relationship between autonomic state and cortical oscillatory activity during NREM sleep, NREM epochs were rank-ordered according to mean heart rate (HR) and divided into 10 equally sized quantile bins within each participant. Mean HR was calculated separately for each NREM epoch using artifact-corrected RR intervals.

For each HR bin, EEG power spectra were estimated using multitaper fast Fourier transform analysis implemented in FieldTrip (48) and averaged across all scalp electrodes. Spectral estimates were obtained using a Hanning taper and subsequently log-transformed before further analyses. Slow-wave activity (SWA) was defined as spectral power between 0.5 and 1.5 Hz, whereas sigma power was defined as spectral power between 12 and 16 Hz.

To quantify HR-related spectral differences, the mean power spectrum of the highest HR bin was compared with that of the lowest HR bin. Percentage power change was calculated separately for each participant as:

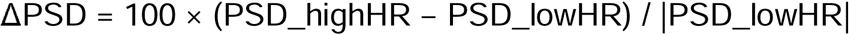

and subsequently averaged across participants.

To characterize the relationship between autonomic state and oscillatory activity, Spearman rank correlations were computed between HR-bin values and both SWA and sigma power. Correlations were calculated using participant-specific HR-bin averages and separately for each scalp electrode. Group-level topographical maps represent the mean correlation coefficient across participants. Participant-wise mean channel correlations were tested against zero using Wilcoxon signed-rank tests.

Frequency-specific differences between high-and low-HR conditions were assessed using nonparametric sign-flipping permutation tests (5,000 permutations). Resulting p-values were corrected for multiple comparisons using the Benjamini–Hochberg false-discovery-rate procedure (62).

### Heart rate variability

High-frequency heart rate variability (HF-HRV) was used as an index of parasympathetic autonomic activity. Artifact-corrected RR intervals obtained from Kubios HRV Scientific (54) were analyzed according to established guidelines (55). Spectral decomposition of HRV was performed in the frequency domain, and high-frequency power (0.15–0.40 Hz) was extracted for each participant. HF-HRV values were averaged across NREM sleep and used as participant-level measures of autonomic state.

To assess the relationship between autonomic regulation and neurocardiac coordination, HF-HRV was subsequently related to SO–spindle coupling, SO–R-peak coupling, and JPAC using sensor-level regression analyses.

### Statistical analyses

All coupling metrics (SO–spindle MI, SO–R-peak EPMI, and JPAC) were computed separately for each electrode. Group-average topographies were visualized using scalp interpolation procedures implemented in FieldTrip (48).

To identify physiological correlates of coupling strength, sensor-level linear regression analyses were performed using HF-HRV, spectral slope, and Lempel–Ziv complexity as predictors. Regression coefficients were estimated separately for each electrode, yielding scalp distributions of regression t-values for each physiological measure. Statistical significance was assessed using cluster-based permutation testing as described below.

Circular phase distributions were assessed using Rayleigh tests and additional circular statistics implemented in CircStat (59). For MI, EPMI, and JPAC, surrogate distributions were generated using 1,000 random permutations while preserving the temporal structure of the original signals.

Topographic consistency across participants was evaluated using the Topographic Consistency Test (63). Sensor-level effects were assessed using nonparametric cluster-based permutation testing (64). Clusters were formed from spatially adjacent electrodes exceeding the predefined statistical threshold, and cluster-level significance was determined using 5,000 random permutations. Unless otherwise stated, significance was defined as two-tailed p < 0.05.

To account for multiple comparisons where appropriate, false-discovery-rate (FDR) correction was applied using the Benjamini–Hochberg procedure (62).

All analyses were performed in MATLAB R2023a (MathWorks, Natick, MA, USA). Data are reported as mean ± standard deviation unless otherwise specified.

## Declaration of AI-Assisted Writing and coding

Generative artificial intelligence (AI) tools (ChatGPT, OpenAI) were used during manuscript preparation to assist with language editing, restructuring of text, formatting, and improvement of readability. The AI tools were not used for data collection, data analysis, statistical testing, interpretation of results, or scientific decision-making.

AI was used to create figure templates and explanatory figures (Fig. 1). Further AI was used to optimize code speed. All scientific content, analyses, conclusions, and final manuscript text were reviewed, verified, and approved by the authors, who take full responsibility for the accuracy and integrity of the work.

## Acknowledgments

We thank Prof. Anita Lüthi for her valuable contributions to our work.

## Author Contributions

Conceptualization: C.A.M., M.P, K.D.F., C.N.

Methodology: C.A.M, K.D.F., T.K., M.dL., V.V., B.F.

Investigation: P.H., E.H., C.L.S. M.A.

Visualization: C.A.M, T.K. K.D.F

Supervision: V.V., D.R., C.N.

Writing—original draft: C.A.M., C.N.

Writing—review & editing: all authors

## Competing Interest Statement

The authors declare they have no competing interests.

## Notes

### Competing Interest Statement

The authors have declared no competing interest.

