## Supplementary material for "Coordinated Brain-Heart Dynamics During Human NREM Sleep": Supp. Material

Prof. Dr. med. Christoph Nissen

Department of Psychiatry, University of Geneva and Geneva University Hospital (HUG)

20 rue de Lausanne

1201 Geneva, Switzerland

-

**Control Analysis**

Cluster-based covariate analyses showed no significant scalp cluster linking NREM heart rate or gender to JPAC. Age, however, was negatively associated with JPAC in a significant centro-frontal cluster. Cluster-mean JPAC decreased with increasing age, r = −0.71, p < 0.001.

For EPMI, no significant clusters were observed for NREM heart rate or gender. Age again showed a significant negative cluster-level association. Cluster-mean EPMI decreased with increasing age, r = −0.61, p < 0.001.

Thus, age—but not resting NREM heart rate or gender—was associated with reduced SO-locked cardiac coupling and reduced tri-modal SO–spindle–R-peak coupling.

Figures

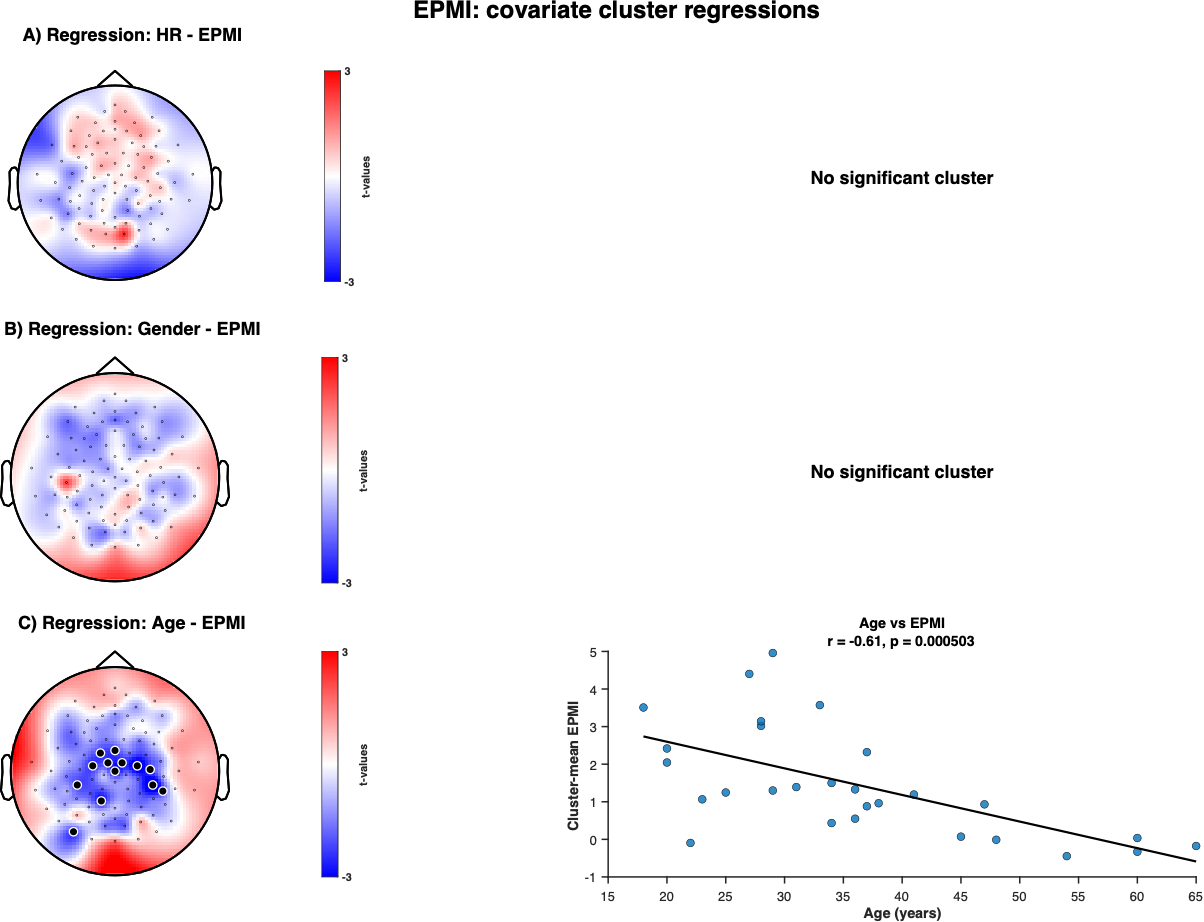

**Fig. S1. Covariate associations with EPMI.**
Topographies show cluster-based regression/contrast t-values for the association between EPMI and A) two-minute mean NREM heart rate, B) gender, and C) age. White circles mark electrodes belonging to the significant cluster. No significant cluster was detected for heart rate or gender. Age showed a significant negative association with EPMI; the scatter plot shows the relationship between age and cluster-mean EPMI (*r* = −0.61, *p* < 0.001).

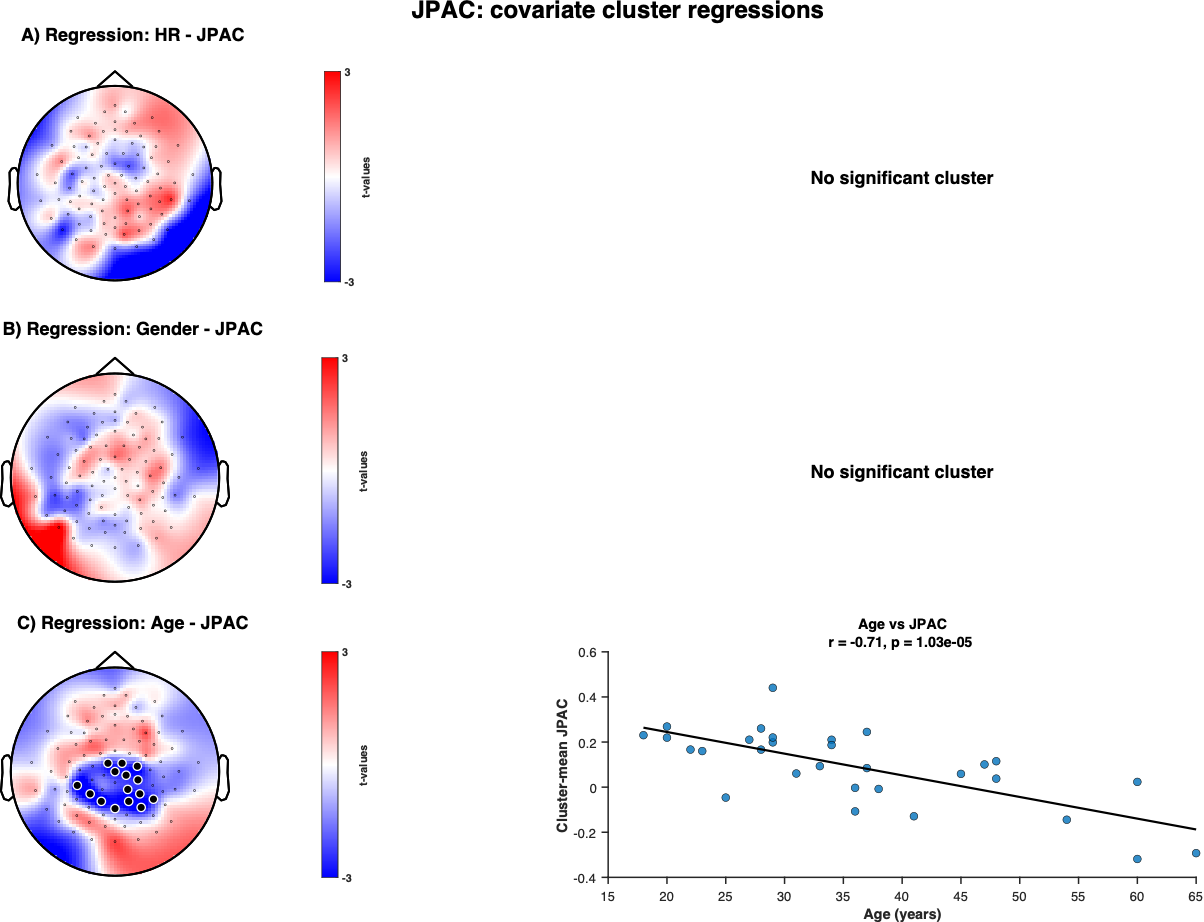

**Fig. S2. Covariate associations with JPAC.**
Topographies show cluster-based regression/contrast t-values for the association between JPAC and A) mean NREM heart rate, B) gender, and C) age. White circles mark electrodes belonging to the significant cluster. No significant cluster was detected for heart rate or gender. Age showed a significant negative association with JPAC; the scatter plot shows the relationship between age and cluster-mean JPAC (*r* = −0.71, *p* < 0.001

Tables

**Table S1. SO and sleep spindle features**

| **Feature** | **Mean ± SEM** |
| --- | --- |
| SO Number of events [N] | 3474 ± 136 |
| SO Duration [s] | 1.374 ± 0.007 |
| SO Amplitude [µV] | 83.36 ± 3.71 |
| SO Frequency [Hz] | 0.768 ± 0.004 |
| Spindle Number of events [N] | 1561.9 ± 106.2 |
| Spindle Amplitude [µV] | 54.56 ± 1.97 |
| Spindle Frequency [Hz] | 12.96 ± 0.05 |
